# Mass spectrometry based immunopeptidomics leads to robust predictions of phosphorylated HLA class I ligands

**DOI:** 10.1101/836189

**Authors:** Marthe Solleder, Philippe Guillaume, Julien Racle, Justine Michaux, HuiSong Pak, Markus Müller, George Coukos, Michal Bassani-Sternberg, David Gfeller

## Abstract

The presentation of peptides on class I human leukocyte antigen (HLA-I) molecules plays a central role in immune recognition of infected or malignant cells. In cancer, non-self HLA-I ligands can arise from many different alterations, including non-synonymous mutations, gene fusion, cancer-specific alternative mRNA splicing or aberrant post-translational modifications. Identifying HLA-I ligands remains a challenging task that requires either heavy experimental work for *in-vivo* identification or optimized bioinformatics tools for accurate predictions. To date, no HLA-I ligand predictor includes post-translational modifications. To fill this gap, we curated phosphorylated HLA-I ligands from several immunopeptidomics studies (including six newly measured samples) covering 72 HLA-I alleles, and retrieved a total of 2,066 unique phosphorylated peptides. We then expanded our motif deconvolution tool to identify precise binding motifs of phosphorylated HLA-I ligands. Our results reveal a clear enrichment of phosphorylated peptides among HLA-C ligands and demonstrate a prevalent role of both HLA-I motifs and kinase motifs on the presentation of phosphorylated peptides. This data further enabled us to develop and validate the first predictor of interactions between HLA-I molecules and phosphorylated peptides.

## Introduction

Human leukocyte antigen class I (HLA-I) molecules mediate cell surface presentation of peptides originating from intracellular protein degradation. Proteins are fragmented by the proteasome into short peptides. These peptides can enter the endoplasmic reticulum through the transporter associated with antigen processing (TAP) protein complex, where they are loaded onto HLA-I molecules and transported to the cell surface (1). HLA-I molecules are encoded by three genes (HLA-A, HLA-B, and HLA-C) and these genes are among the most polymorphic of the human genome, resulting in currently more than 17,000 different alleles (2). HLA-I molecules have specific binding motifs and different alleles typically bind distinct sets of peptides (3). Foreign or altered-self HLA-I ligands can be recognized by CD8+ T cells and induce an immune response to eliminate infected or malignant cells. These non-self HLA-I ligands can have various origins, such as viral proteins or genetically or post-translationally modified proteins in cancer. Identification of peptides presented HLA-I molecules is labor intensive since it relies either on challenging immunopeptidomics experiments, or predictions of HLA-I ligands followed by experimental validation. Currently, many optimized algorithms are available for predicting unmodified HLA-I ligands (4–7), but none of them include specifically post-translational modifications.

Multiple studies have identified post-translational modifications (PTMs) showing aberrant behavior in cancer (8–11), resulting for instance in abnormal cellular signaling, one of the hallmarks of cancer (12). Phosphorylation of serine, threonine, and tyrosine is one of the most frequent and best studied PTMs (13), and is carried out by different types of protein kinases, consisting mainly of serine/threonine- and tyrosine-specific protein kinases. There are over 500 known kinases in the human genome (14, 15) and different phosphorylation motifs, such as [pS/pT]P for CDK1 or MAPK1 or Rxx[pS/pT] for PKA or PKB, characterize individual kinases. It has been shown that aberrant phosphorylation can occur in cancer cells due to a deregulated balance between phosphorylation and dephosphorylation events (16), thereby altering key signaling pathways and processes within cells. A recent study estimated that phosphorylation-related single nucleotide variants are present in approximately 90% of tumor genomes (17) and predicted that 29% of these variants affect signaling pathways. Furthermore, phosphosite-specific signature analysis showed to be able to identify dysregulation of phosphorylation-regulated pathways in cancer (18).

Peptides with phosphorylated residues can be processed by the antigen presentation pathway, bind to HLA-I molecules, and be presented on the cell surface (19–30). Several studies reported that phosphorylated peptides could induce immune responses through T cell recognition. For instance, T cells were shown to recognize phosphorylated peptides presented on primary tumors and normal tissues and kill tumor cell lines (26, 31). Studies have reported a clear preference for the phosphorylation at position 4 on HLA-I ligands and revealed an increased presence of arginine at P1 as well as an enrichment of proline after the phosphosite caused by proline-dependent kinases (20, 21, 24, 26–30, 32, 33). However, so far, only a handful of phosphorylated HLA-I ligands were determined for a small number of HLA-I alleles and no method is available to specifically predict their binding. As a result, predictions of HLA-I interactions with phosphorylated ligands are performed without including the modified amino acid (either by using the unmodified version of the residue or by substituting it with ‘X’), which is likely sub-optimal since no information about phosphorylation is included in the training set of the predictors.

To fill this gap, we first measured the immunopeptidome of 6 new samples and reprocessed existing immunopeptidomics raw data for 55 other samples, in order to search for phosphorylated HLA-I ligands. We complemented these data with a small subset of HLA-I restricted phosphorylated peptides identified previously by mass spectrometry (MS), some of which came from phospho-enrichment protocols, and curated a large dataset of 2,066 unique phosphorylated HLA-I ligands experimentally determined by MS. This enabled us to accurately determine phosphorylated motifs for 22 of the most frequent HLA-I alleles and revealed clear discrepancies among alleles in terms of propensity to bind phosphorylated peptides. In particular, we observed a much higher frequency of phosphorylated ligands for HLA-C alleles. We further analyzed several properties of phosphorylated HLA-I ligands and performed binding assays to validate and interpret these results. Using these data, we then developed the first predictor of phosphorylated HLA-I ligands.

## Experimental Procedures

### Preparation of HLA class I peptide samples

Several tissue samples, 3993, 4052-BA, 3989-HT, OE37-1N, OVZW-1P and OXVD-09, were provided by the biobank of the Center of Experimental Therapies at the CHUV after informed consent of the participants was obtained following requirements of the institutional review board (Ethics Commission, CHUV). HLA typing of the cell lines was experimentally determined prior to this work and is provided for all samples in this study. 2–5 biological replicates per tissue were processed using our previously described protocol (34). Briefly, tissues were homogenized on ice in lysis buffer with Ultra Turrax homogenizer (IKA, Staufen, Germany) for 10 seconds at maximum speed, and then incubated on ice for 1 hr. Lysis buffer contained 0.25% sodium deoxycholate (Sigma-Aldrich), 0.2 mM iodoacetamide (Sigma-Aldrich), 1 mM EDTA, 1:200 Protease Inhibitors Cocktail (Sigma, Missouri, USA), 1 mM Phenylmethylsulfonylfluoride (Roche, Mannheim, Germany), 1% octyl-beta-D glucopyranoside (Sigma) in PBS. Subsequently, 20 min centrifugation for clearance (table-top centrifuge, Eppendorf Centrifuge 5430R, Schönenbuch, Switzerland) was performed at 4°C at 14200 rpm. Immuno-affinity purification through Protein-A Sepharose beads covalently bound to W6-32 antibodies was performed in a format of 96-well single-use micro plate with 10 µm polypropylene membranes (SeaHorse Bioscience). The plates were then washed 4 times with 2mL 150 mM NaCl and 20 mM Tris HCl (buffer A), 4 times with 2mL 400 mM NaCl and 20 mM Tris Hcl, further 4 times 2mL buffer A and final twice with with 20 mM Tris HCl, pH 8. HLA molecules and peptides were eluted with 1% trifluoroacetic acid (TFA, Merck, Darmstadt, Switzerland) directly into Sep-Pak tC18 100 mg Sorbent 96-well plates (Waters, Massachusetts, USA) pre-conditioned with 80% acetonitrile (ACN) in 0.1% TFA and with 0.1% TFA only. Wells were washed twice with 0.1% TFA and then the peptides were eluted with 28% ACN in 0.1% TFA. Peptides were dried using vacuum centrifugation (Eppendorf Concentrator Plus, Schönenbuch, Switzerland) and were resuspended in a final volume of 12 uL 0.1% TFA. 3 uL of these peptides were used for each MS run.

### Mass spectrometry analysis of HLA class I peptides

HLA peptides were separated by a nanoflow HPLC (Proxeon Biosystems, Thermo Fisher Scientific, Odense) on 50 cm long column (75 μm inner diameter) self-packed with ReproSil-Pur C18-AQ 1.9 μm resin (Dr. Maisch GmbH, Ammerbuch-Entringen, Germany) in buffer A (0.5% formic acid) coupled on-line to a Q Exactive HFX mass spectrometers (Thermo Fisher Scientific, Bremen) with a nanoelectrospray ion source (Proxeon Biosystems). HLA-I peptides were eluted with a linear gradient of 2–30% buffer B (80% ACN and 0.5% formic acid) at a flow rate of 250 nl/min over 125 min. MS spectra were acquired from m/z = 300-1650 in the Orbitrap with a resolution of 60’000 (m/z = 200) and ion accumulation time of 80 ms. The auto gain control was set to 3e6 ions. MS/MS spectra were acquired on 10 most abundant precursor ions with a resolution of 15,000 (m/z = 200), ion accumulation time of 120 ms and an isolation window of 1.2 m/z. The auto gain control was set to 2e5 ions. Dynamic exclusion to 20 s and a normalized collision energy of 27 was used for fragmentation. The peptide match option was disabled. No fragmentation was performed in case of assigned precursor ion charge states of four and above.

### Identification of HLA class I peptides

We employed the MaxQuant platform (35) version 1.5.5.1 to search the MS peak lists against a fasta file containing the human UniProt database containing 42,170 entries including isoforms (March 2017) and a list of 247 frequently observed contaminants. Peptides with a length between 8 and 15 amino acids were allowed. The second peptide identification option in Andromeda was enabled. The enzyme specificity was set as unspecific and FDR of 5% was required for peptides and no protein FDR was set. As a large score difference to the second best match (delta score) is important for identification of phosphorylated peptides (36), the delta score was set to a minimum of 10 for both modified and unmodified peptides. The initial allowed mass deviation of the precursor ion was set to 6 ppm and the maximum fragment mass deviation was set to 20 ppm. Methionine oxidation (15.994915 Da), N-terminal acetylation (42.010565 Da) and phosphorylation (79.9663304 Da) on serine, threonine and tyrosine were set as variable modifications.

### Experimental Design and Statistical Rationale

In addition to the six novel samples mentioned above, 38 MS samples from published immunopeptidomics studies were reanalyzed together, 209 raw files in total (6, 28, 34, 37, 38). For each sample, at least two technical replicates of raw MS files were included. In a separate run, the MaxQuant platform was employed with the same parameters on 85 MS raw files of 17 monoallelic samples with five technical replicates of raw MS files for each sample (39), and the data was similarly filtered to obtain peptide spectrum matches with high confidence.

### Curation of immunopeptidomics HLA-I MS datasets

We filtered the list of identified phosphorylated HLA-I peptides listed in the MaxQuant MSMS output table by removing reverse hits and peptides matching contaminants. To maintain peptide spectrum matches with high confidence, the list was further filtered by restricting the identification score ≥70, and the localization probabilities to ≥0.75. Only unique modified and unmodified sequences were further analyzed.

Additionally, data from various publications (20, 21, 23–27, 29, 30, 32, 33, 40–43) was added to our dataset, using from each sample both, known phosphorylated as well as unmodified HLA-I binders if available. Identified phosphorylated HLA-I peptides from enrichments studies (20, 23, 24, 27, 30) were included in the determination of phosphorylated HLA-I binding motifs and the training of the predictor, but not in the comparison of the fraction of phosphorylated ligands for different HLA-I molecules.

### HLA-I Motif Deconvolution for identification of HLA-I binding motifs

To determine allelic restriction among HLA-I ligands found by MS, including phosphorylated peptides, we expanded our motif deconvolution tool MixMHCp (6, 44) to allow for additional non-standard amino acids. Briefly speaking, the motif deconvolution method infers with Expectation-Maximization algorithm K different position weight matrices that optimally model the list of peptides. In this extended version of the motif deconvolution algorithm, phosphorylated residues are treated as additional amino acids, leading to an alphabet of size 23 (i.e., position weight matrices of size 9×23, instead of 9×20 as described in our previous manuscript (44)). Finally, motifs were assigned to their respective HLA-I allele using the approach described in (37) and all assignments were manually curated. Of note, MixMHCp also contains a flat motif to which peptides that do not match any of the motifs inferred by the algorithm are assigned. The command-line script to run the motif deconvolution (MixMHCp2.1) can be obtained at https://github.com/GfellerLab/MixMHCp.

### Visualization of HLA-I Phosphorylated Motifs

Binding motifs of HLA-I alleles were visualized by sequence logos. The sequence logos were generated by modifying the R package ggseqlogo (45) in a way to include sequences with modified amino acids. Purple letters were used to visualize phosphorylated residues in sequence logos of HLA-I binding motifs. Phosphorylated motifs for HLA-I alleles with more than 22 phosphorylated ligands are displayed in Fig. 1. The modified version of ggseqlogo to plot sequence logos including modified residues is provided at https://github.com/GfellerLab/ggseqlogo.

**Figure 1:**
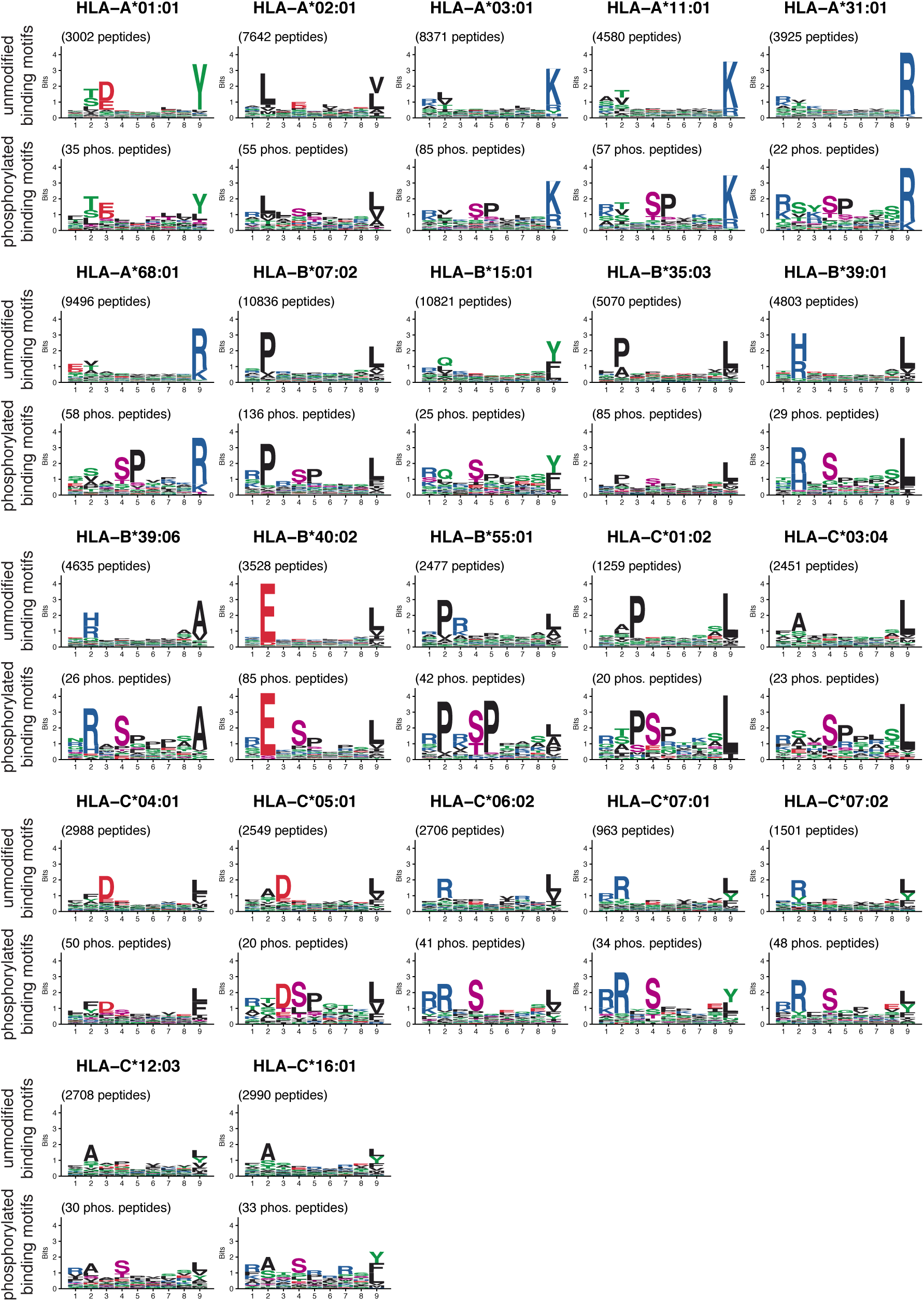
Overview of 9-mer HLA-I binding motifs of unmodified (top) and phosphorylated (bottom) ligands for HLA-I alleles with at least 20 phosphorylated ligands (9-mers) determined in this work. Phosphorylated residues are shown in purple.

### Analysis of Phosphorylated HLA-I Ligands

After assigning unmodified and phosphorylated peptides to alleles through motif deconvolution, peptides were pooled from all samples for each allele and merged into a unique set of peptides per allele. Phosphorylated and unmodified binding motifs for each HLA-I allele were built. The overall frequency of phosphorylated peptides per allele was analyzed by computing the fraction of phosphorylated peptides among all discovered peptides per allele from any length and for each peptide length ranging from 8 to 12 amino acids separately. To identify potential structural differences between binding regions of alleles with high and low frequency of phosphorylated peptides, alleles were split into two groups based on the median frequency in HLA-A, -B, or -C alleles, respectively. HLA-I binding sites were analyzed by (1) selecting positions of the binding regions that show interaction with the peptide in the 3D structure (28 positions in total) and (2) computing the Euclidean distances for these selected positions of the binding regions between the two groups. For each group of alleles, the block of binding site sequences was transformed into position weight matrices (*M_ij_*) with *i* = 1, …, 20 and *j* = 1, … , 28 by calculating the frequency of each amino acid *i* at each position *j*. For each position *j*, the Euclidean distance between the columns of the matrices (*M*(^1^) and *M*(^2^)) was computed as:

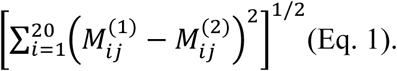

Sequence logos were used to visualize the ten most different positions in each comparison for binding sites of alleles with high frequency versus alleles with low frequency of phosphorylated peptides.

For each allele in the underlying dataset, we calculated how many unmodified ligands contained phosphosites from the phosphoproteome (46). In particular, phosphosites positioned at P4 in unmodified HLA-I 9-mers were counted for all alleles with more than 50 unmodified ligands. The frequency of known phosphosites per allele was computed as the fraction of detected phosphosites at P4 within the unmodified 9-mer HLA-I ligands. The correlation between phosphorylated HLA-I ligands per allele and the amount of detected phosphosites within unmodified HLA-I ligands was measured using the Pearson correlation coefficient.

The length distribution ranging from 8- to 12-mers was computed for phosphorylated and unmodified HLA-I ligands per allele and error bars show the variability across alleles.

The distribution of phosphorylated amino acids (pS, pT, and pY) in the human phospho-proteome was obtained from (46) and compared to the one observed in the whole phosphorylated immunopeptidome and separately for each length 8 to 12. P-values were calculated by t-tests. Furthermore, among all unique phosphorylated HLA-I ligands, we measured how often each position in any 8-to 12-mer was phosphorylated.

To test if proline enrichment exists in our dataset, the proline frequency in phosphorylated and unmodified HLA-I ligands was analyzed. Firstly, the frequency of proline occurring next to a phosphorylated residue was measured in all phosphorylated peptides per allele, for all alleles with at least 5 phosphorylated peptides. Secondly, the overall proline frequency in unmodified HLA-I peptides at non-anchor positions (3 to 8 for HLA-I 9-mers) was extracted allele-wise. As a third measurement, the proline frequency in the human proteome (UniProt as of October 2017) was also included as a comparative means in the analysis of proline enrichment. P-values were computed by t-tests between the different groups of data.

To compute the enrichment in arginine at P1, for each allele with at least 5 phosphorylated HLA-I ligands the occurrence of arginine at P1 was calculated among all phosphorylated peptides with a phosphorylated P4 (phosphorylated serine, threonine or tyrosine), which is the most frequent phosphorylated position within 9-mers. These values were compared to (1) the frequency of arginine at P1 in unmodified HLA-I ligands with serine, threonine, or tyrosine at P4 and (2) the overall frequency of arginine in the human proteome (UniProt as of October 2017). P-values comparing the different measurements were computed with t-tests.

Binding motifs of different kinases were determined with the Phospho.ELM dataset (47) and sequence logos of 3 positions upstream and 3 positions downstream of the phosphosite were visualized with the modified version of ggseqlogo.

### Experimental Testing of HLA-I–Phosphorylated Ligand Binding

Experimental testing of HLA-I ligands was performed as described before (6), consisting of refolding assays, followed by ELISA assays. ELISA absorbance signals were used to define binding stabilities of phosphorylated and unmodified versions of several peptides and several alleles. Two replicates per experiment were performed and negative controls correspond to experiments performed in the absence of a peptide. Measured absorbance of the binding assays were normalized by t=0h of the positive controls. Half-lives were computed as ln(*2*) /*k_off_*. The background signal (i.e. measurements from the negative controls) was removed from the measured ELISA absorbance values and *k_off_* was determined through fitting exponential curves to absorbance values.

Mutation of position 69 for arginine to alanine in the heavy chain of HLA-C*06:02 was done by site-directed mutagenesis by overlap extension using the polymerase chain reaction (PCR). Two PCR products are obtained from the HLA-C*06:02 BSP coding sequence using as forward primer 5’-GATATACATATGTGCTCCCACTCCATGAGG-3’ (primer A) and reverse primer containing the mutation (in bold and underlined) 5’-CACTCGGTCAGCCTGTGCCTG**<u>GGC</u>**CTTGTACTTCTGTGTCTCCCG-3’ (primer B) and second PCR with forward primer containing the mutation (in bold and underlined) 5’-CGGGAGACACAGAAGTACAAG**<u>GCC</u>**CAGGCACAGGCTGACCGAGTG-3’ (primer C) and reverse primer 5’-GGCCGCAAGCTTTTAGTGCCATTCGATTTTCTGAGC-3’ (primer D). The two PCR products are mixed together in a third PCR with primer A and D. The coding sequence was cloned between *NdeI* and *Hind III* sites in plasmid pET-23a. Expression of mutated R69A HLA-C*06:02 was performed by using the *Escherichia coli* strain BL21(DE3)(pLysE).

### Predictor and Cross Validation

For each HLA-I allele with at least 20 phosphorylated HLA-I 9-mer ligands, position weight matrices (PWM) were built. PWMs are then used to calculate a peptide score for each peptide (*X*_1_, … *X*_*L*_):

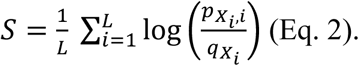

The peptide score describes for a peptide with which frequency each amino acids occurs at its position in the binding motif of the allele. *L* corresponds to the length of the predicted peptide, *px_i_,i* is the PWM entry at position *i* for amino acid *X_i_*, and *qx_i_* is a background frequency. Here, average frequencies of each amino acid within the human phospho-proteome are used as background frequencies. For each allele, peptides are ranked according to their score to identify most likely binders.

Prior to calculating the peptide score *S*, a pseudocount is added to the PWM, as described in (48). This is done to prevent zero occurrence of any amino acid at any position in the PWM, which may arise especially for small training datasets. The pseudocount for phosphorylated PWMs (PWMs of size 9×23 for 9-mers) is based on the work on the BLOSUM62 alignment score (49). The transition probabilities from the original BLOSUM62 were used for unmodified amino acids. BLOSUM62 was then expanded to include the three phosphorylated residues (phosphorylated serine, phosphorylated threonine, and phosphorylated tyrosine), based on the BLOSUM62 transition probabilities of unmodified serine, threonine and tyrosine. The phosphorylated-BLOSUM62 was extended by (1) transition probabilities from each phosphorylated amino acid to any of the three phosphorylated as well as any of the 20 unmodified amino acid, and (2) transition probabilities from any unmodified amino acid into each of the three phosphorylated residues. In more details, for phosphorylated residues *p* ∈ (*s*, *t*, *y*) and unmodified residues *U* ∈ (*A*, *C*, *D*, ⋯, *Y*) BLOSUM62 was extended into phospho-BLOSUM62 in the following way: transitions *b* from any phosphorylated amino acid *p*_1_ to any other phosphorylated amino acid *p*_2_ were defined as

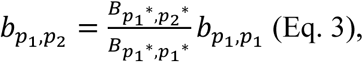

with *p*^∗^ denoting the corresponding unmodified amino acid of phosphorylated amino acid *p* and *B* corresponding to transition probabilities from the original BLOSUM62. Furthermore,

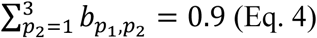

was defined to strengthen the transition probability of a phosphorylated amino acid *p*_1_ to stay phosphorylated and to have only little probability to transform into any unmodified residue (sum of transition probability into unmodified residues being 0.1). The transition *b* from phosphorylated residue *p*_1_ to any unmodified amino acid *U* was set to

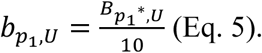

Transitions of the unmodified counterpart of *p*_1_ to *U* from the original BLOSUM62 were used to reflect the relationship between two amino acids also in the transitions from the phosphorylated version of the amino acid into all unmodified residues, but only contributing with a reduced weight to the total transitions for a phosphorylated amino acid (factor 10 is used to reduce the row sum of the original BLOSUM62 from 1 to 0.1, considering Eq. 4). For any unmodified amino acid *U* the transition probability into a phosphorylated residue *p* was defined as

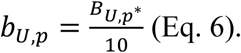

Reduced transition probabilities were defined to provide a low transition probability from unmodified into phosphorylated amino acids, yet modeling a similar proportion for the transitions into phosphorylated versions of the amino acids like the transitions into unmodified serine, threonine, and tyrosine. For each unmodified amino acids, a row-wise normalization over the transitions to all unmodified and all phosphorylated amino acids was performed.

Various versions of the predictor with different training data compositions were developed to find the best prediction method. One predictor was trained only on phosphorylated peptides. A second predictor included all unmodified HLA-I ligands of the allele in addition to the phosphorylated peptides in the training set. In addition, to avoid over-fitting of the predictor, if the unmodified version of a phosphorylated peptide in the testing data was present among unmodified HLA-I ligands, it was removed from the training dataset. A third predictor was trained exclusively on unmodified HLA-I binders. In this version of the predictor every phosphorylated residue in the prediction data was treated like an unmodified amino acid, since no information about phosphorylated residue occurrence was given in the PWM. Furthermore, the performance of the different versions of the predictor is also benchmarked against NetMHCpan4.0 (5). NetMHCpan was carried out on the same prediction data replacing the phosphorylated residues by ‘X’ and peptides were ranked according to their Rank score.

To validate the performance of the above-described predictors with varying training data, a 5-fold cross validation was performed on all alleles with more than 20 phosphorylated HLA-I peptides of length 9 and run 100 times. For each allele the set of phosphorylated peptides was randomly divided into five groups, four were used as training data for the predictor and the remaining one was used as positive testing data. In addition, four times the amount of positive peptides was added as negative data to the testing data. Negative peptides were randomly selected from a pool of all known phosphosites of the human proteome, excluding phosphosites at anchor positions 2 and 9. The Area Under the ROC Curve (AUC), AUC0.1 as well as precision of the top 20% of predicted phosphorylated peptides (corresponding to recall as four fifth of the prediction data was negative data) were used to measure the performance of the different prediction models (50) and shown as a mean over all 100 runs averaged over all alleles and for each allele separately. P-values comparing the results of the cross validation were computed using Wilcoxon signed-rank test.

The code of the predictor of phosphorylated HLA-I ligands is available at https://github.com/GfellerLab/PhosMHCpred.

## Results

### Identification of phosphorylated HLA-I ligands uncovers phosphorylated HLA-I binding motifs

We collected data from six new samples and curated 55 publicly available immunopeptidomics studies (6, 28, 34, 37–39) comprising both pooled and mono-allelic datasets. None of these MS studies were performed with phospho-enrichment protocols. Raw MS data were reprocessed allowing for phosphorylation on serine, threonine and tyrosine as variable modifications (see Experimental Procedures). To gain sensitivity while maintaining peptide spectrum matches with relatively higher confidence that is typically achieved with the more conservative false discovery rate (FDR) of 1%, we applied FDR of 5% and considered peptides identified with Andromeda search engine score ≥70, score difference to the second best peptide spectrum match (delta score) ≥10 and localization probability ≥0.75. This resulted in 2,190 unique phosphorylated peptides in total for all 61 samples. Fig. S1A shows the distribution of Andromeda search engine peptide spectrum match score and delta score for phosphorylated peptides with different localization probabilities. To determine allelic restriction, we expanded our motif deconvolution algorithm MixMHCp (6, 44) to consider both the phosphorylated and unmodified peptides in each sample (see Experimental Procedures). MixMHCp removes potentially wrongly identified peptides that do not match the inferred motifs by assigning them to a so-called flat motif (6). 1,841 unique phosphorylated peptides (84.1%) were assigned to HLA-I motifs following this deconvolution step, while the remaining phosphorylated peptides were assigned to the flat motif and excluded from downstream analyses. More than 31% of peptides identified with delta score <20 were assigned to the flat motif compared to only ∼12% of the relatively more reliable identifications of peptides with delta score ≥20 (see Fig. S1B). We then compared different properties (i.e., peptide length, position of the phosphosite and frequency of the different phosphorylated residues) between phosphorylated HLA-I ligands with delta score ≥10 or ≥20. Peptides assigned to HLA-I alleles by MixMHCp displayed very similar properties across different choices of delta score thresholds (Fig. S1C). Reversely, peptides assigned to the flat motif showed a dramatically different behavior suggesting that the filtering with MixMHCp efficiently removes several wrongly identified peptides (Fig. S1C). Similar properties were also observed between phosphorylated peptides assigned to HLA-I motifs when using FDRs of 5% or 1% (Fig. S1D). Finally binding motifs of phosphorylated peptides were robust to different choices of thresholds on delta scores and FDRs (see Fig. S2).

We further curated phosphorylated HLA-I ligands with known allelic restriction reported in earlier studies (20, 21, 23–27, 29, 30, 32, 33, 40–43), including phosphorylated peptides from five samples analyzed with phospho-enrichment protocols (20, 23, 24, 27, 30). In the final step, we restricted HLA-I peptides to length 8-12 for further analysis. Altogether, a total number of 2,066 unique phosphorylated peptide sequences were retrieved, representing 2,585 unique HLA-I–phosphorylated peptide interactions with 72 different HLA-I alleles (20 HLA-A, 30 HLA-B, 21 HLA-C alleles and 1 HLA-G allele). 740 of the 2,585 (28.63%) unique HLA-I–phosphorylated peptide interactions had been reported in previous studies. Phosphorylation occurred mostly once per peptide (97.77%) and very few multiple phosphorylated peptides were found (2.23% double phosphorylated peptides, see also Fig. S3A). Among the 2,066 phosphosites, 800 (38.7%) were not observed in phosphosite databases (dbPAF, phosphoELM, and phosphositePLUS (51–53)), which is in line with what had been previously reported (28). Comparison of binding motifs of all phosphorylated peptides with binding motifs excluding unknown phosphosites showed that these were very similar (see Fig. S2, column 1 and 4). 171 unique phosphorylated HLA-I ligands (8.3%) were only found in the five phosphorylation-enriched samples from previous studies (20, 23, 24, 27, 30) and not in any of the other unenriched samples included in our work. This demonstrates that many phosphorylated HLA-I ligands can be detected in samples that are not specifically enriched in phosphorylation residues, as already shown in previous studies (28). ∼ 30% of the unique interactions between HLA-I alleles and phosphorylated peptides were also detected in their unmodified version. A GO enrichment analysis of source proteins of all HLA-I ligands showed very similar proteins of origin for phosphorylated and unmodified peptides (Fig. S3B).

22 HLA-I alleles had more than 20 unique phosphorylated 9-mer ligands and their phosphorylated and unmodified binding motifs are shown in Fig. 1. These 22 alleles were later used to develop the predictor for HLA-I–phosphorylated peptide interactions and have a population frequency of 91.76% on average worldwide (99.41% in Europe).

### Phosphorylated Peptides are enriched among HLA-C ligands

We analyzed the fraction of phosphorylated HLA-I ligands for alleles derived from each gene (HLA-A, HLA-B, and HLA-C). Phosphorylation-enriched samples (20, 23, 24, 27, 30) were not considered in this analysis to prevent biases in the estimation of the frequency of phosphorylated HLA-I ligands. For all peptide lengths combined, we observed variability across different alleles and a significant enrichment of phosphorylated peptides among HLA-C alleles compared to HLA-A and HLA-B (Fig. 2A and see Fig. S4A for separate lengths). To show that this could not be ascribed to the peptides from the monoalleleic HLA-C samples (39), we performed the same analysis without these samples, and obtained similar results (Fig. S4B, left panel). Furthermore, we could see that these results were also robust when only analyzing phosphorylated peptides identified with delta score ≥20 (Fig. S4B, right panel). To investigate why specific alleles, and especially HLA-C alleles, show a higher fraction of phosphorylated peptides in our dataset, we measured the binding stability differences (i.e., half-life ratio) between phosphorylated and unmodified peptides for several HLA-A, -B, and -C alleles (HLA-A*01:01, HLA-A*25:01, HLA-B*07:02, HLA-B*18:01, HLA-C*06:02, and HLA-C*07:02) with both high and low phosphorylated peptide frequency in immunopeptidomics data (arrows in Fig. 2A). Overall, the results indicate significantly higher half-life ratio for HLA-C alleles compared to HLA-A or HLA-B alleles (Fig. 2B). However, the selected HLA-A and HLA-B alleles showed opposite binding preference compared to what would have been expected from the frequency of phosphorylated peptides in the immunopeptidome (Fig. 2B, compare HLA-A*01:01 and HLA-A*25:01, HLA-B*07:02 and HLA-B*18:01).

**Figure 2:**
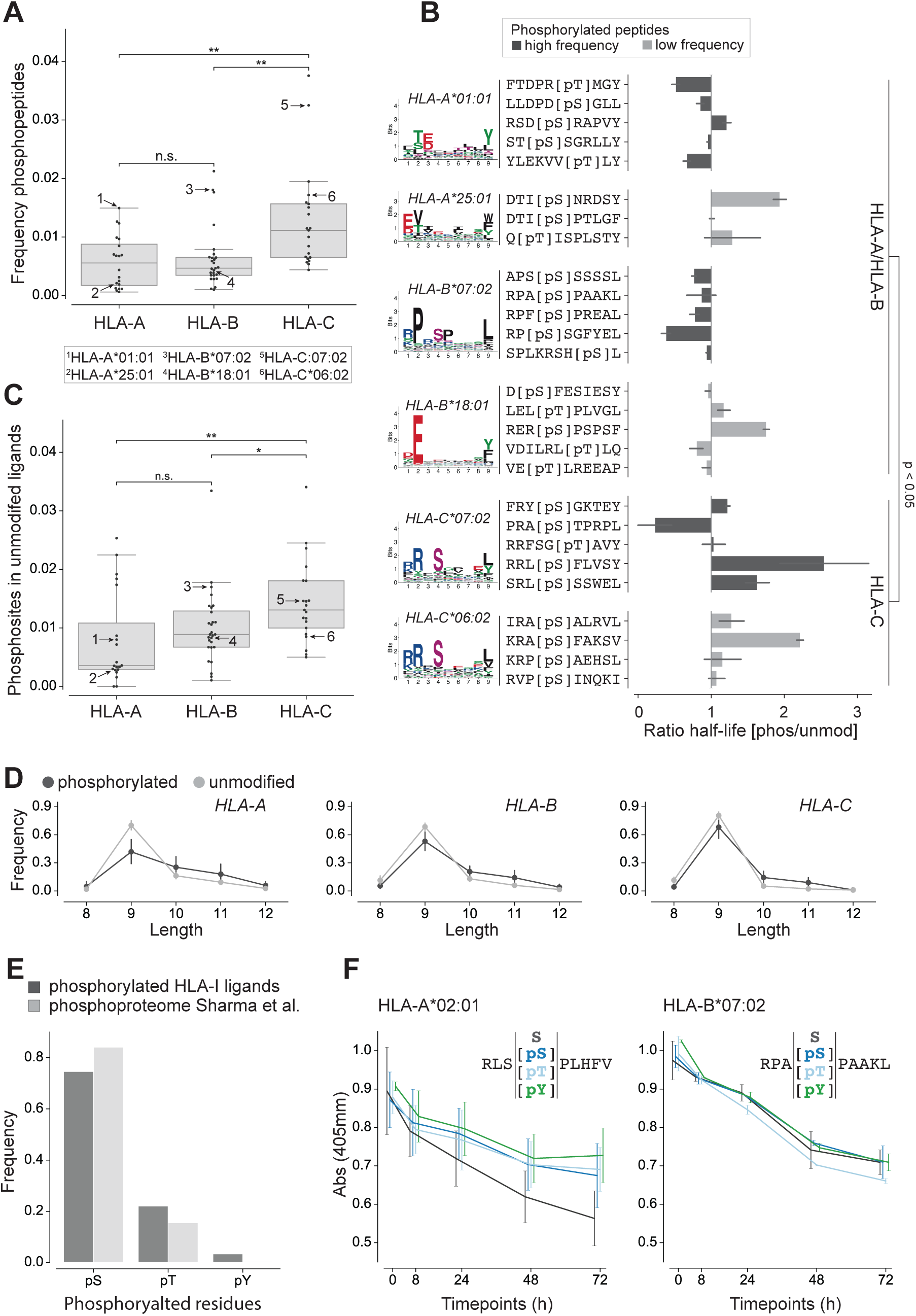
Analysis of phosphorylated peptides across HLA-I alleles. **(A)** Frequency of phosphorylated peptides per HLA-A, -B, and -C alleles for peptides of any length. Numbers in the plot indicate alleles tested in panel B. **(B)** Ratio of half-lives between the phosphorylated (pS/pT) and the unmodified (S/T) peptides for several alleles. The colors of the bars correspond to alleles with high and low frequency of phosphorylated peptides in A. For HLA-A*01:01, HLA-B*07:02, HLA-C*06:02 and HLA-C*07:02 phosphorylated HLA-I binding motifs are shown, for HLA-A*25:01 and HLA-B*18:01 binding motifs of unmodified HLA-I ligands are given because too few phosphorylated peptides were observed in MS data for these alleles. **(C)** Fraction of unmodified HLA-I 9-mer ligands containing a phosphosite at P4 for HLA-A, -B, and -C alleles. Arrows indicate the same alleles as in panel A. **(D)** Length distribution of phosphorylated and unmodified ligands of HLA-A, HLA-B, and HLA-C alleles. **(E)** Frequency of the different phosphorylated residues within phosphorylated HLA-I ligands of length 8 to 12 and within the human phosphoproteome (46). **(F)** Dissociation assay (absorbance from ELISA) for unmodified and phosphorylated peptides (with phosphorylated serine, phosphorylated threonine, and phosphorylated tyrosine). (*: p <= 0.05, **: p <= 0.01)

HLA-C molecules are characterized by the specific presence of R at position 69 (A/T in HLA-A or -B), which may interact with phosphorylated residues (Fig. S4C). However, R69A mutation did not show a sharper decrease in binding affinity for phosphorylated ligands (Fig. S4D), suggesting that this residue is not responsible for the preference of HLA-C alleles for phosphorylated peptides. Moreover, comparison of binding sites between alleles with high and low fractions of phosphorylated ligands did not suggest clear differences that could favor the binding of phosphorylated peptides (Fig. S4E).

The lack of correlation between stability measurements and fraction of phosphorylated peptides in MS data within alleles from the same gene and the lack of molecular features explaining the variability observed across alleles (Fig. 2A-B and Fig. S4A-E) led us to hypothesize that this variability may be related to a better compatibility of specific HLA-I motifs with phosphorylation motifs. To explore this hypothesis, we analyzed the unmodified ligands of each allele in our dataset by checking for the occurrence of human phosphosites from the phosphoproteome (46). The results showed that the frequency of unmodified ligands containing known phosphosites at P4 is on average higher in HLA-C than in HLA-A or HLA-B alleles (Fig. 2C). We could furthermore detect a significant positive correlation between the amount of phosphosites of the human phosphoproteome found within unmodified ligands and the frequency of phosphorylated ligands detected per allele (Fig. S4F). These results support the hypothesis that HLA-I alleles that preferentially bind phosphorylated peptides (especially HLA-C alleles) have motifs that are better suited to bind peptides coming from known phosphosites. (Fig. 2A and Fig. S4A). Overall, our analysis shows large variability in the fraction of phosphorylated ligands across alleles and suggest that some of this variability comes from a better compatibility between the HLA-I motifs of specific alleles and phosphorylation motifs.

When we compared the fraction of phosphorylated peptides for different lengths, we could observe that longer peptides are enriched with phosphosites (Fig. 2D and Fig. S4A). Furthermore, phosphorylated residues (phosphorylated serine, threonine, and tyrosine) were observed at similar frequencies as in the human phospho-proteome (46) (Fig. 2E), and binding assays indicated no difference in binding stability for different phosphorylated residues (Fig. 2F).

### Phosphorylated HLA-I Ligands show a Preference for Phosphosites at P4 which does not only results from higher binding stability

Fig. 3A shows the distribution of phosphorylated positions for peptides of lengths 8 to 12 in our phosphorylated immunopeptidome. Similar to what has been shown before (26, 28, 30, 32, 33), we could detect a clear preference for phosphorylation at P4 in all lengths and an additional preference for P6 in 10- and 12-mer phosphorylated HLA-I ligands. To explore the biochemical reason for this preference at P4, the binding of 9-mer peptides with phosphorylated serine at non-anchor positions 3 to 8 was tested. As expected, HLA-A*02:01 and HLA-B*07:02 binding peptides with a phosphorylated P4 bind more stably compared to peptides with phosphorylated positions 3 and 5 to 7. This trend was also observed for a peptide that was originally found in the MS data with a phosphorylated P3 (third peptide of HLA-B*07:02 in the panel). Furthermore and less expectedly, the results demonstrated that phosphorylation at P8 also shows an increased binding compared to phosphorylated positions 3 and 5 to 7 (Fig. 3B), especially for ligands tested for HLA-B*07:02.

**Figure 3:**
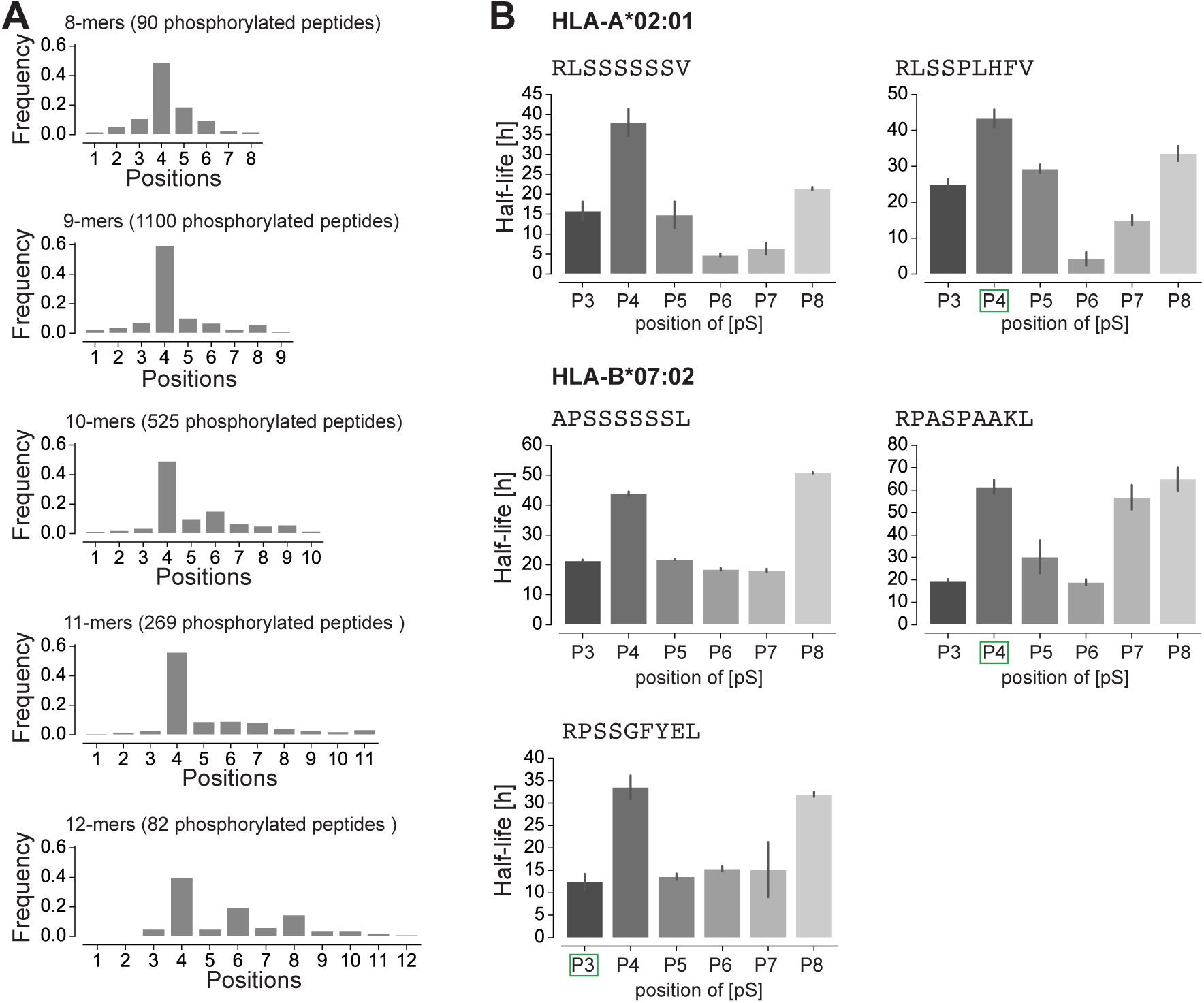
Phosphorylated positions in HLA-I ligands. (**A)** Distribution of the position of phosphorylated residues in phosphorylated HLA-I ligands of lengths 8 to 12. (**B)** Half-lives of HLA-I ligands for peptides with positions 3 to 8 substituted by phosphorylated serine. Green squares mark the position of the phosphosite (phosphoserine) for peptides found in MS data. Lack of green square indicates one unmodified peptide observed in MS data (APSSSSSSL) or one synthetic peptide (RLSSSSSSV) used in this *in vitro* assay.

### Proline Adjacent to the Phosphorylated Residue and Arginine at P1 are a Result of Kinase Motifs

Previous studies reported proline enrichment next to phosphorylated residues in phosphorylated HLA-I ligands as a consequence of the [pS/pT]P phosphorylation motif (20, 26–28, 30, 33). Our analysis with a much larger allelic coverage confirmed these results. In particular, we observed a significantly higher frequency of proline next to phosphorylated residues in HLA-I ligands compared to unmodified HLA-I ligands or to the human proteome (see Experimental Procedures and Fig. 4A and Fig. S5A). To support the hypothesis that the higher frequency of proline next to phosphorylated residues in HLA-I ligands reflects kinase phosphorylation motifs (Fig. 4B), we performed binding assays for three alleles (HLA-A*02:01, HLA-A*11:01, HLA-B*07:02) and four peptides with or without a phosphorylated residue at P4 and with or without proline at P5. The results of these binding assays show that proline or alanine at P5 did not change the binding stability, both for phosphorylated and unmodified peptides, consistent with the hypothesis that the proline enrichment is mainly due to kinase phosphorylation motifs (Fig. 4C).

**Figure 4:**
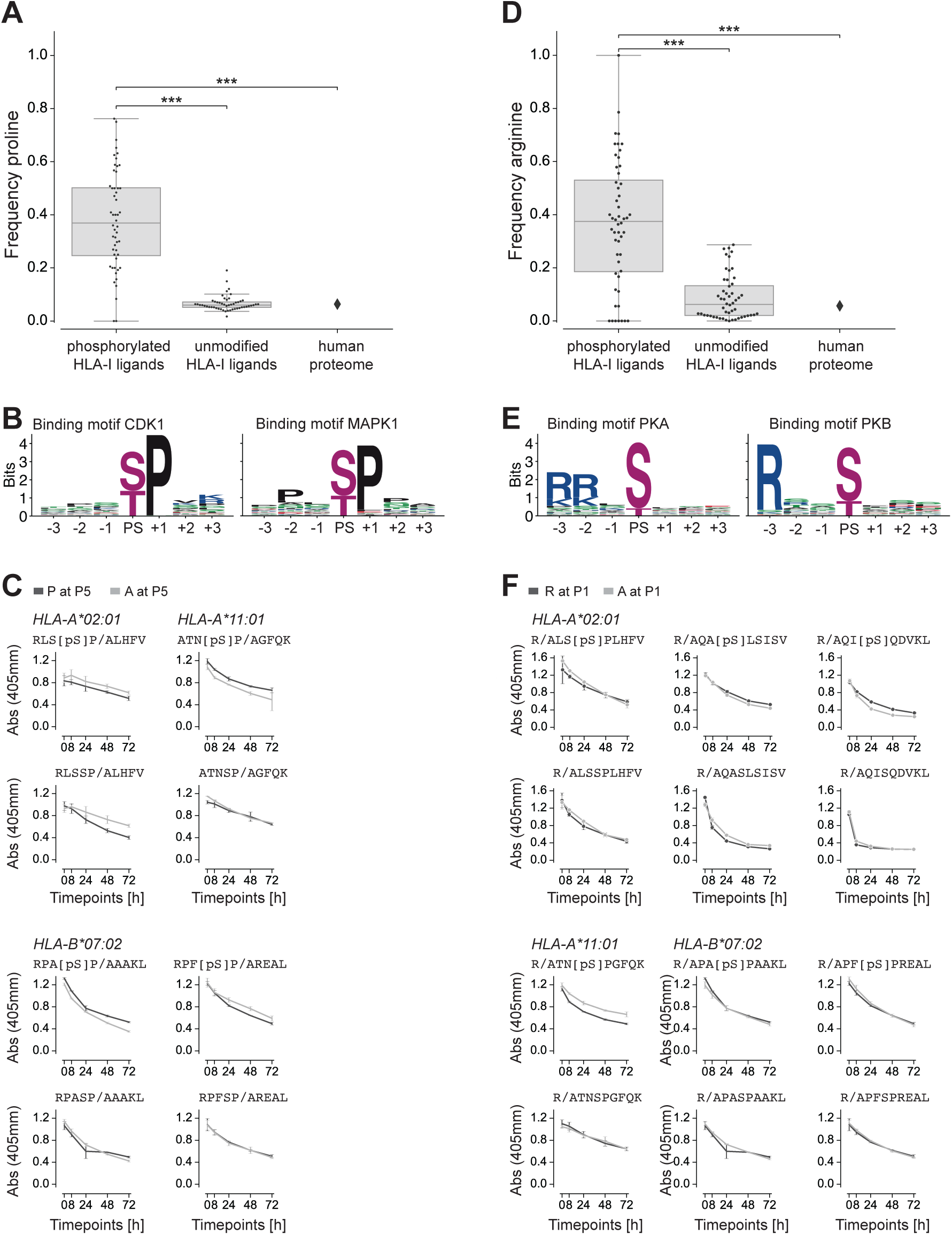
Proline and arginine enrichment in phosphorylated HLA-I ligands. (**A)** Frequency of proline next to phosphorylated residues in phosphorylated HLA-I ligands, proline at non-anchor positions in unmodified HLA-I ligands, and proline frequency in the human proteome. (**B)** Kinase binding motifs for kinases CDK1 and MAPK1, three positions up- and three positions downstream of the phosphosite (PS). (**C)** Dissociation of peptides with proline or alanine next to phosphorylated serine (top) and next to unmodified serine in unmodified versions of the peptides (bottom). (**D)** Frequency of arginine at P1 in phosphorylated HLA-I ligands, in unmodified HLA-I ligands, and in the human proteome. **(E)** Kinase binding motifs for kinases PKA and PKB, three positions up- and downstream of the phosphosite (PS). (**F)** Dissociation of peptides with arginine at P1 compared to peptides with alanine at P1 for both the phosphorylated (top) and unmodified (bottom) versions of the peptides. (***: p <= 0.001)

Previous studies have shown that some phosphorylated HLA-I ligands show a preference for basic amino acids at P1 (20, 24, 26, 28, 30, 32, 33). For several alleles, we could see the same trend in our data when comparing phosphorylated and unmodified binding motifs (Fig. 1, e.g. HLA-B07:02 or HLA-C06:02). Comparisons between the frequency of arginine at P1 in phosphorylated HLA-I ligands with the frequency of arginine at P1 in unmodified HLA-I ligands and in the human proteome confirmed that the enrichment is statistically significant (see Experimental Procedures and Fig. 4D and Fig. S5B). We then asked whether this is due to enhanced binding of phosphorylated HLA-I ligands in the presence of R at P1 or to a signature of the known Rxx[pS/pT] phosphorylation motifs of protein kinases (see examples for PKA and PKB in Fig. 4E). To this end, we measured the binding stability of six different peptides with or without phosphorylated serine at P4 and with or without arginine at P1. For multiple alleles and peptides, the peptides with R at P1 have a similar binding compared with the peptides with A at P1, for both the phosphorylated and unmodified versions (Fig. 4F), suggesting that the preference for R at P1 does not affect the binding of phosphorylated ligands, but rather results from the phosphorylation motifs of specific kinases. Table S1 shows phosphorylated HLA-I ligands that correspond to known phosphosites from the phosphoELM database (52) and were identified to be phosphorylated by kinases CDK1, PKA and PKB.

### Training predictors on HLA-I phospho-peptidomes improves predictions of phosphorylated HLA-I ligands

We then used our large curation of eluted phosphorylated peptides to train a predictor of phosphorylated HLA-I ligands. In particular, we explored different alternatives consisting of training the predictor (1) combining both the phosphorylated and non-phosphorylated peptides, (2) considering only unmodified peptides or (3) considering only phosphorylated peptides (see Experimental Procedures). For validation, we focused on the 9-mer peptides and performed a 5-fold cross validation on the 22 alleles with at least 20 phosphorylated peptides. Phosphorylated ligands per allele were divided into training and testing dataset and negative peptides were added by randomly selected peptides from the human phosphosite reference database (see Experimental Procedures). Fig. 5A-C show the area under the receiver operating characteristics (ROC) curve (AUC), AUC0.1, and the precision for the top 20% of predicted phosphorylated peptides, respectively, of the cross validations for each version of training data for each allele separately (see Fig. S6A-C for average values over all 22 alleles). The results indicate that training the predictor with a combination of phosphorylated and unmodified HLA-I ligands performs best (1^st^ bar, Fig. 5A-C and Fig. S6A-C). When comparing the AUC values of the predictor trained with a combined dataset (1^st^ bar) and the predictor trained only with unmodified peptides (2^nd^ bar), we can see that training only on unmodified HLA-I ligands and ignoring modification on a residue is also not as good as training on a combined dataset for the prediction of HLA-I–phosphorylated peptide interactions. Finally, we observed that our predictor trained on a combined dataset of phosphorylated and unmodified HLA-I ligands outperforms the state-of-the-art predictor NetMHCpan4.0 (4^th^ bar in Fig. 5A-C and Fig. S6A-C) (phosphorylated residues were substituted with ‘X’ since NetMHCpan cannot take phosphorylated amino acids as input). This further confirms that including phosphorylated residues within the training dataset of HLA-I ligand predictors improves their accuracy for predicting phosphorylated peptides.

**Figure 5:**
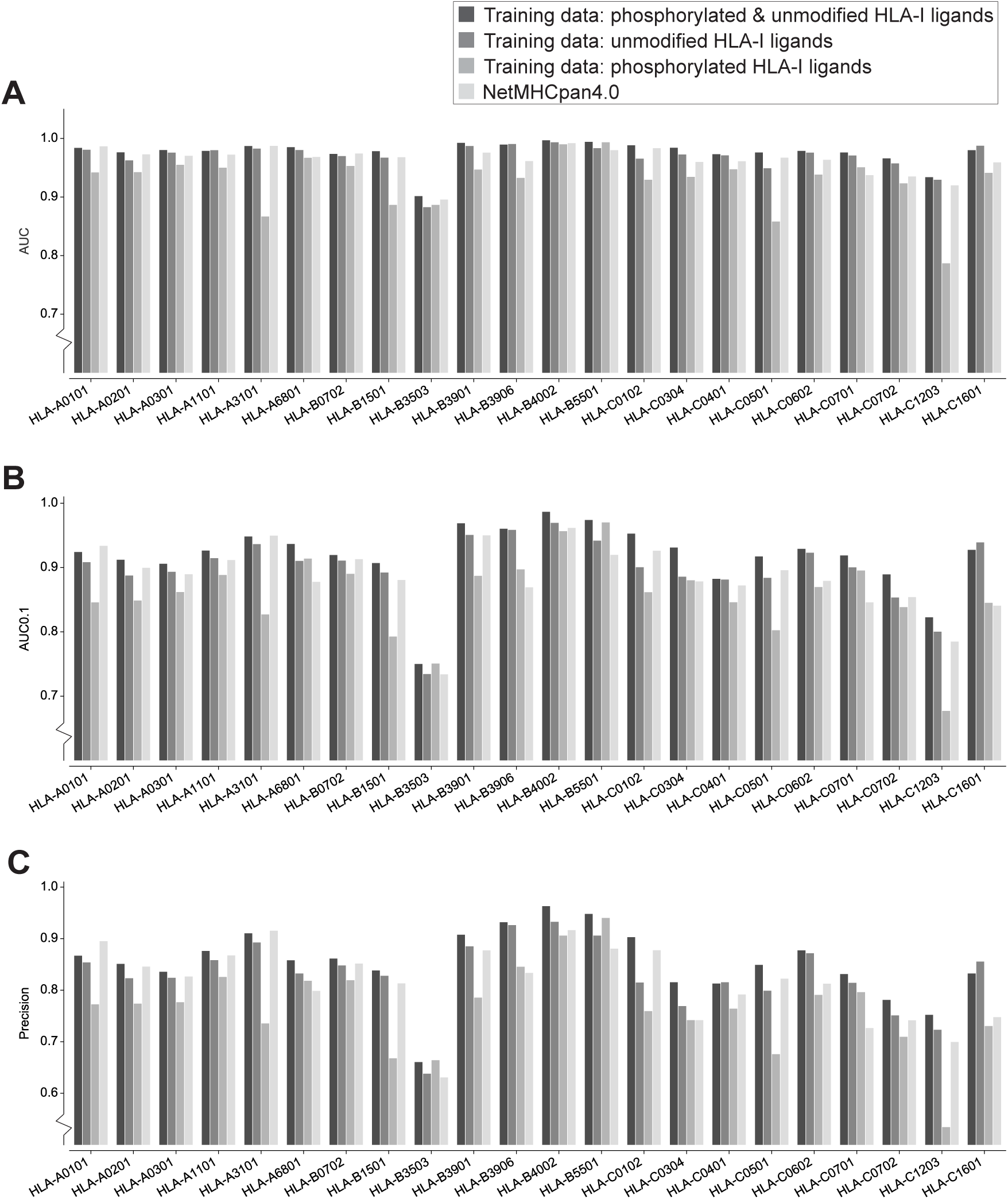
Cross validation of the predictor for each HLA-I alleles with more than 20 phosphorylated 9-mer peptides. **(A)** AUC values for phosphorylated HLA-I 9-mer peptides when trained on both phosphorylated and unmodified ligands (1^st^ bar), trained only on unmodified ligands (2^nd^ bar, treating phosphorylated residues as their unmodified counterpart), or when trained only on phosphorylated HLA-I ligands (3^rd^ bar). For comparison, AUC values are also shown when using NetMHCpan4.0 and replacing phosphorylated residues by ‘X’ in the input (4^th^ bar). **(B)** Results of the 5-fold cross validation measured by AUC0.1. **(C)** Precision measured for the top 20% of the predicted peptides (equivalent to recall of the prediction data).

To test the robustness of our predictor to the presence of some level of wrongly identified phosphorylated peptides, we included 5% phosphorylated decoy peptides in the training data. We did not observe significant changes in predictions, which demonstrates that our predictor can tolerate some level of contaminants or wrongly identified peptides (Fig. S6D). To exclude potential batch effects in our data, we trained the predictor on the newly curated data and tested it on previously reported HLA-I-restricted phosphorylated peptides. This was done for two alleles (HLA-A*02:01 and HLA-B*07:02) with sufficient previously reported HLA-I restricted phosphorylated peptides. Fig. S6E shows that the results from these predictions are similar to the case where the training and testing data are selected from the whole pool of peptides (i.e., without distinguishing our newly curated data from previously reported data). A similar analysis was performed for ligands from phospho-enriched samples. Fig. S6F shows that prediction of enriched samples trained on data from non-enriched samples performs equally well than randomly selected training and testing data of the same size. Finally, a saturation analysis with different amount of phosphorylated ligands in the training data was performed for HLA-A*02:01 (Fig. S6G), showing that the choice of 20 phosphorylated peptides is already providing good prediction accuracy.

## Discussion

Aberrant phosphorylation is frequent in malignant cells. However, prediction of the presentation of phosphorylated peptides on HLA-I molecules has been poorly explored, mainly due to the lack of training data. Here, we curated phosphorylated HLA-I ligands across many immunopeptidomics studies to investigate molecular properties of interactions between phosphorylated peptides and HLA-I molecules and develop the first predictor for HLA-I interactions with phosphorylated peptides.

Our unsupervised approach to assign allelic restriction and infer binding motifs based on motif deconvolution (6, 37, 44) is especially appropriate for phosphorylated peptides since it does not require a priori information on their interactions with HLA-I alleles. In addition, it enabled us to use relatively permissive thresholds and subsequently filter potentially wrongly identified peptides that did not match the inferred motifs. As expected, the resulting phosphorylated motifs show similarity with those derived from unmodified peptides, especially at anchor positions (second and last positions for most HLA-I alleles).

Our results outlined a clear preference for phosphorylated peptides to bind to HLA-C alleles compared to HLA-A and -B alleles (Fig. 2A). Yet, differences in the binding site, in particular R69 in the HLA heavy chain, do not appear to determine the preference for phosphorylated peptides of HLA-C alleles. Previous work (30) identified the interaction between R62 in HLA-B*40 and the phosphate moiety of the phosphorylated HLA-I ligand to support the binding of the phosphorylated peptide to this allele. However, all HLA-C as well as most HLA-B alleles in our dataset, including those with low fractions of phosphorylated ligands, contain arginine at position 62 (Fig. S4G). This suggests that, at least for the alleles studied in this work, arginine at position 62 does not necessarily favor the binding of phosphorylated HLA-I ligands. However, the analysis of unmodified ligands showed a higher fraction of human phosphosites in HLA-C ligands, suggesting that phosphosites fit the binding motifs of HLA-C alleles on average better than those of HLA-A and HLA-B alleles (Fig. 2C). This hypothesis is further supported by the correlation between the number of detected phosphosites per allele and the number of phosphorylated HLA-I ligands (Fig. S4F). This could explain, at least partly, the higher fraction of phosphorylated HLA-C ligands observed in immunopeptidomics data (Fig. 2A).

Our results demonstrated a clear preference for phosphorylated residues at P4 in phosphorylated HLA-I ligands, confirming previous observations (28, 30, 32). However, binding assays for HLA-B*07:02 pointed out that peptides with phosphorylation at P8 show a similar binding stability as peptides with phosphorylation at P4 (Fig. 3B). 9-mer binding motifs of alleles shown in Fig. 1 indicate that proline at P9 is not favorable for binding. This can explain why peptides with phosphorylation at P8 are less often observed in MS data, since the common kinase motif with proline next to the phosphorylated residue is not compatible with the binding motifs of HLA-I alleles. This suggests that the clear specificity of phosphorylation at P4 is due to (1) better binding of these phosphorylated peptides and (2) incompatibility of kinase motifs for phosphorylated peptides with phosphorylation at P8.

Consistent with what has been shown in previous studies (20, 21, 24, 26, 28, 30, 32, 33), we observed a clear preference for arginine at P1 for several alleles (Fig. 4D and Fig. S5B). Basic residues at P1 were observed to interact with the negatively charged phosphorylated residue at P4, and were suggested to improve the general stability of the phosphorylated peptide with the HLA-I molecule through this intramolecular bond (21, 30). However, when we tested the binding of phosphorylated and unmodified peptides with both arginine or alanine at P1, we observed very similar binding stability, suggesting that arginine does not specifically strengthen the binding of phosphorylated peptides in these alleles. Of course, we cannot exclude that the intramolecular bridge may be present and enhance binding of phosphorylated peptides to other alleles. Moreover some alleles show preference for arginine at P1 for both phosphorylated and unmodified peptides, which could explain why arginine at P1 was reported to enhance binding of phosphorylated peptides (21, 30). Yet, the analysis of different kinase binding motifs indicates that many serine/threonine-kinases have a binding motif with arginine three positions upstream of the phosphorylated residue (Fig. 4E). This, together with the results of binding assays in Fig. 4F, provides a more likely explanation for the enrichment in arginine at P1 observed in Fig. 1 and Fig. 4D. Of note, arginine at P1 is not observed in all alleles (e.g., HLA-A*68:01 where P1 serves as an additional anchor residue, see Fig. 1).

Our results demonstrated that a combined training dataset of phosphorylated and unmodified peptides outperforms training that is based solely on unmodified peptides. Hence, we can conclude that without phosphorylated peptides in the training data the predictor lacks important information about the phosphorylated residue (especially the preference for the phosphorylated residue at P4)

Overall, our analysis of phosphorylated immunopeptidomes shows that the presentation of phosphorylated peptides on HLA-I molecules is governed by a combination of HLA-I binding motifs (specificity mainly at P2 and PΩ), intrinsic HLA-I binding properties of phosphorylated peptides (specificity at P4) and kinase motifs (specificity at P1 and P_phospho_+1). Our ability to integrate these different features into a robust predictor of phosphorylated HLA-I ligands explains the improvement over existing tools and provides a rationale for training on both unmodified and phosphorylated HLA-I ligands. In this work, we have leveraged existing immunopeptidomics data for the identification of a large collection of phosphorylated HLA-I ligands, mostly originating from un-enriched samples. To develop the predictor for as many alleles as possible, we applied relatively permissive filters and thresholds for interpretation the MS immunopeptidomic dataset because motif deconvolution can filter many potentially wrongly identified peptides (6) (see also a related strategy for general immunopeptidomics experiments (54)) and our computational approach can handle some level of false positives. This is in contrast to other application of MS based immunopeptidomics studies aimed at directly identifying novel epitopes, where highly confidence peptide identification is crucial. We anticipate that this predictor will facilitate the identification of phosphorylated T cell epitopes for researchers that do not have access to high-quality but expensive MS immunopeptidomics technology and will foster future research on their role in immune recognition of infected or malignant cells.

## Acknowledgment

We are thankful to Stefan Stevanovic and Moreno Di Marco for sharing with us the raw MS data from (39). We are thankful to Julien Schmidt and the Protein and Peptide Chemistry Facility from UNIL for synthesizing the peptides.

## Data availability

The MaxQuant output tables, the list of phosphorylated HLA-I ligands, and the mass spectrometry immunopeptidomics raw files will be made available upon publication.

## Notes

**Declaration of Interest:** The authors declare no potential conflicts of interest.

