## Supplemental Information for "Mass spectrometry based immunopeptidomics leads to robust predictions of phosphorylated HLA class I ligands"

### Supplementary Figures

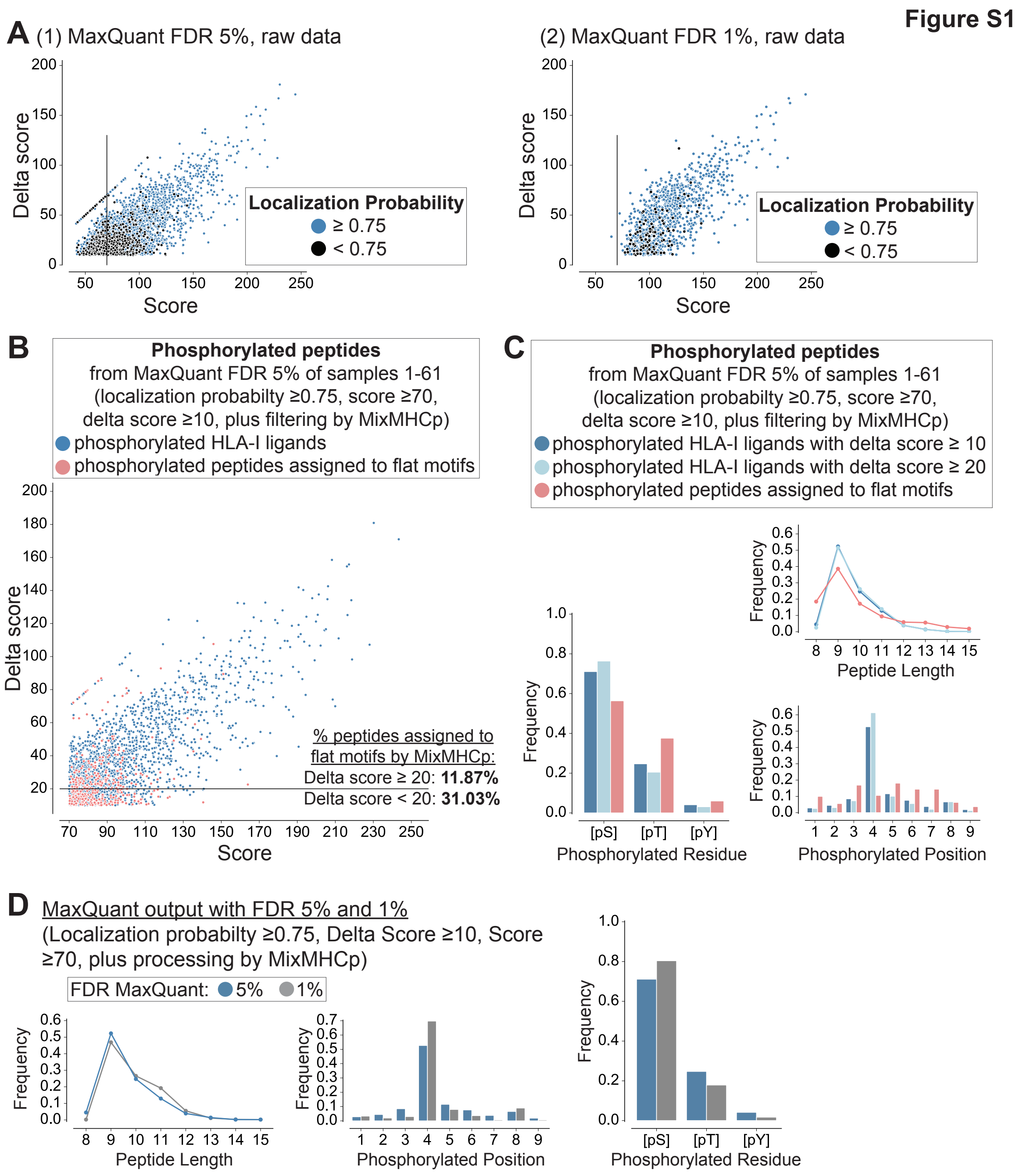


**Figure S1: (A)** Distribution of Andromeda search engine scores vs. score differences to the second best peptide spectrum match (delta scores) for MaxQuant search with FDR of 5% (left) and with FDR of 1% (right). Blue dots mark phosphorylated peptides with localization probability of ≥0.75, black dots mark phosphorylated peptides with localization probability <0.75. **(B)** Distribution of Andromeda search engine scores vs. delta scores for all peptides assigned to HLA-I alleles (blue dots) and the flat motif (red dots) in the motif deconvolution. **(C)** Distribution of phosphorylated residues ([pS/pT/pY]), peptide lengths, and phosphorylated positions in 9-mers for phosphorylated peptides assigned to HLA-I alleles with delta scores ≥10 (blue) and delta scores ≥20 (cyan), and for peptides assigned to the flat motif by MixMHCp (red). **(D)** Distribution of phosphorylated residues ([pS/pT/pY]), peptide lengths, and phosphorylated positions in 9-mers for phosphorylated peptides assigned to HLA-I alleles with FDR of 5% (blue) and FDR of 1% (grey).

**Figure S2**

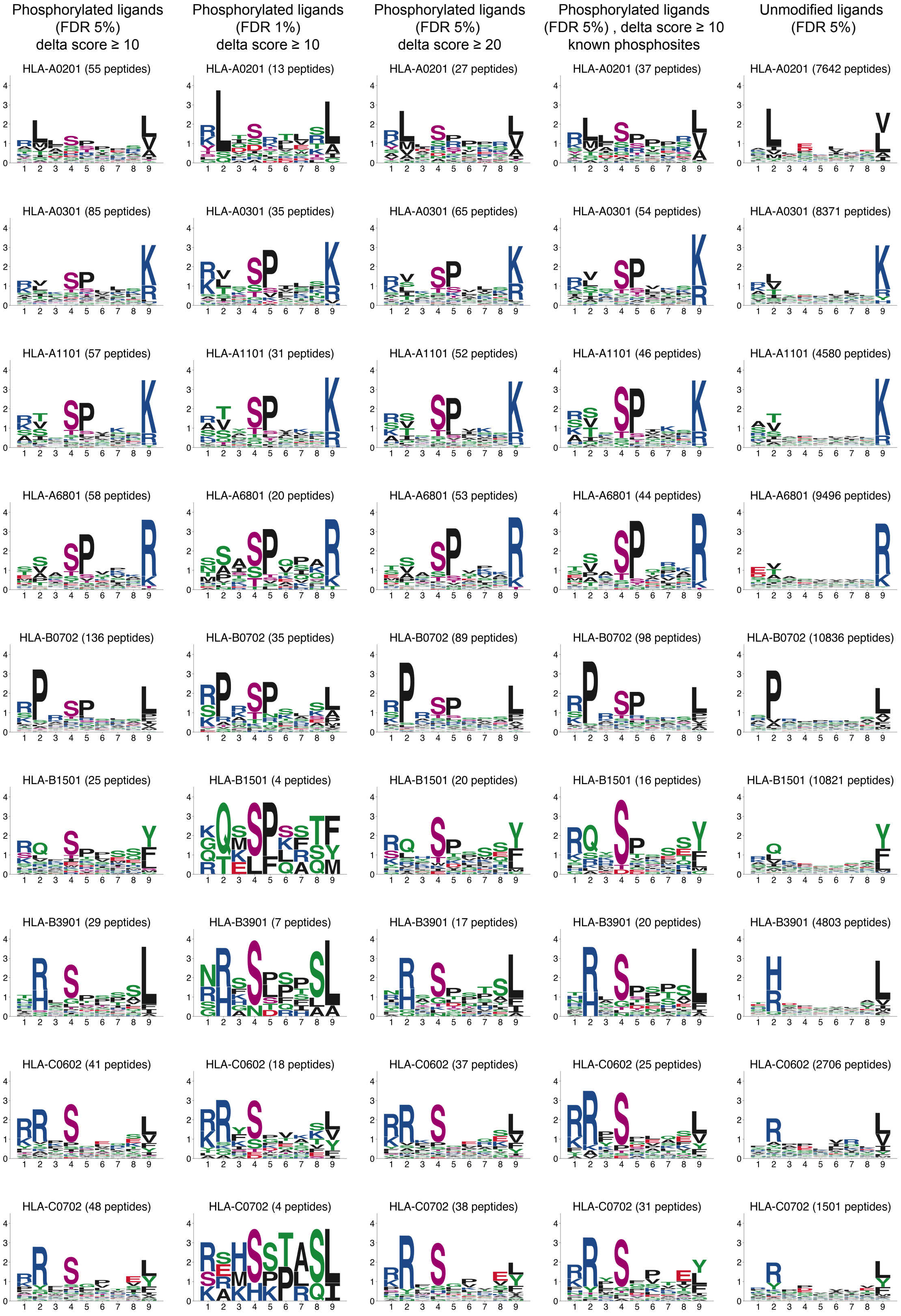


**Figure S2:** Comparison of binding motifs for different choices of parameters used to search the MS data. Column 1 shows the motifs based on phosphorylated peptides with FDR of 5% (same data as in Figure 1). Column 2 shows the motifs based on phosphorylated peptides with FDR of 1%. Column 3 shows the motifs based on phosphorylated peptides with delta score ≥20. Column 4 shows the motifs based only on phosphorylated peptides containing known phosphosites. Column 5 shows the motifs based on unmodified peptides.

**Figure S3**

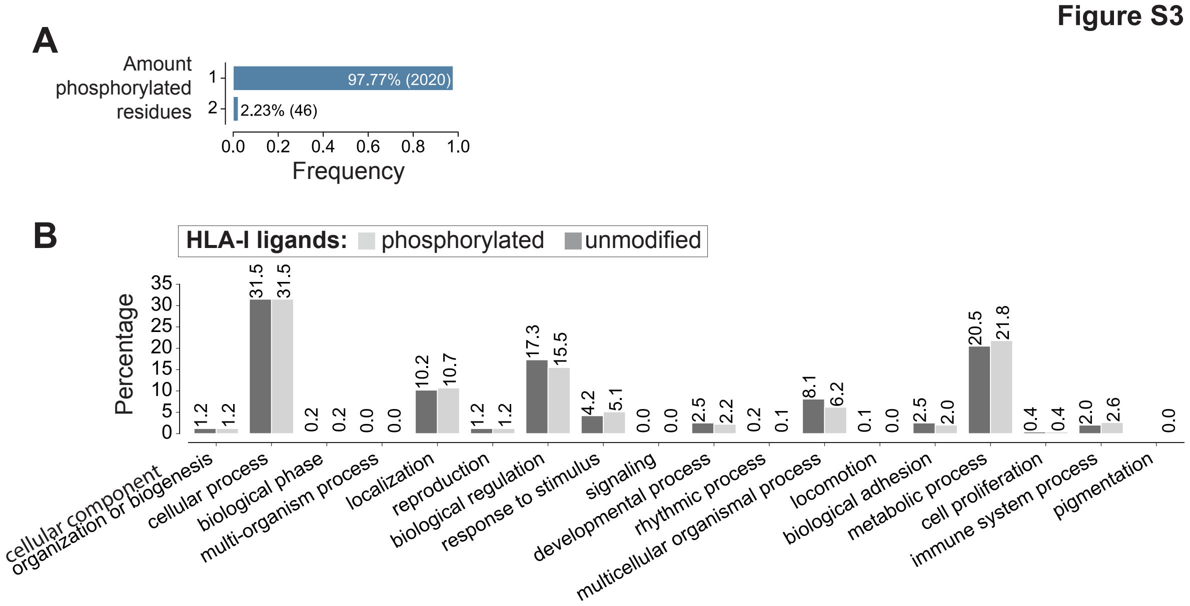


**Figure S3: (A)** Amount of phosphorylated residues per HLA-I ligand. **(B)** GO enrichment analysis with biological process classification for the source proteins of all phosphorylated and unmodified HLA-I ligands (performed with Panther tool [1]).

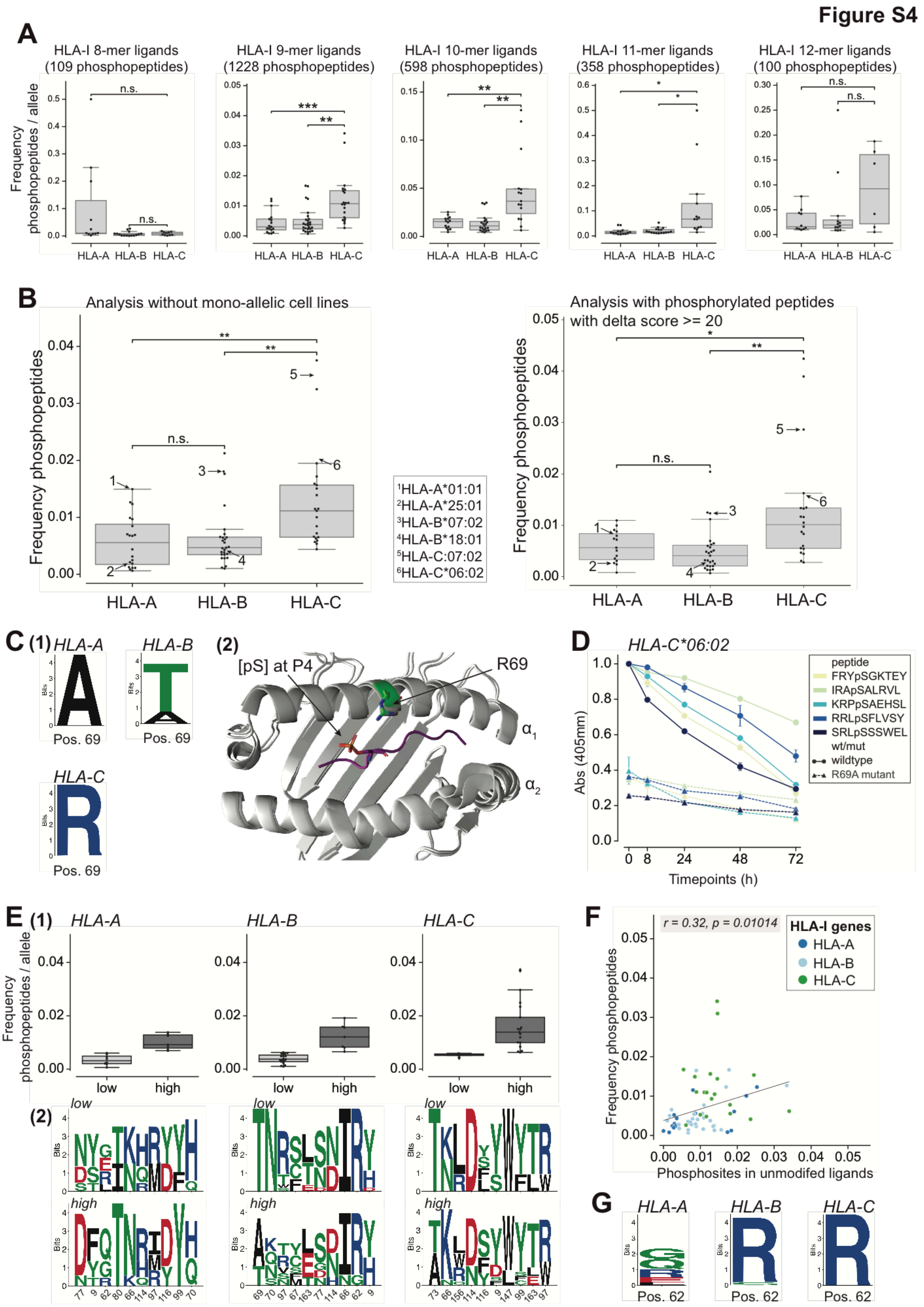


**Figure S4:** Analysis of phosphorylated peptides per HLA-I allele. **(A)** Frequency of detected phosphorylated peptides per allele for different lengths. **(B)** Frequency of phosphorylated peptides per HLA-A, -B, and -C alleles for peptides of any length without monoallelic HLA-C samples (left panel) and for HLA-I ligands with delta score ≥20 (right panel). Arrows indicate same alleles as in Fig. 2A and C and correspond to alleles tested in Fig. 2E. (**C**) Sequence logos of position 69 in HLA-A, -B, and -C alleles (1). Crystal structure of HLA-C*06:02 (PDB code: 5w67) superimposed to HLA-A*02:01 (PDB code: 4nnx) with phosphorylated ligand RQA[pS]LSISV. R69 of HLA-C*06:02 is shown in green (2). **(D)** Dissociation assays of phosphorylated peptides with HLA-C*06:02 wt (R at P69) and mutated (R69A) alleles. **(E)** Frequency of phosphorylated peptides per allele grouped into high and low alleles. Sequence logos of the ten most different positions of the HLA-I binding site (measured by Euclidean distance between the groups with high and low alleles) for high and low alleles of HLA-A, HLA-B and HLA-C separately. **(F)** Correlation of phosphosites from the human proteome found in unmodified ligands and the frequency of phosphorylated HLA-I ligands per allele. **(G)** Sequence logos of position 62 in HLA-A, HLA-B, and HLA-C alleles. (*: $p \le 0.05$; **: $p \le 0.01$; ***: $p \le 0.001$)

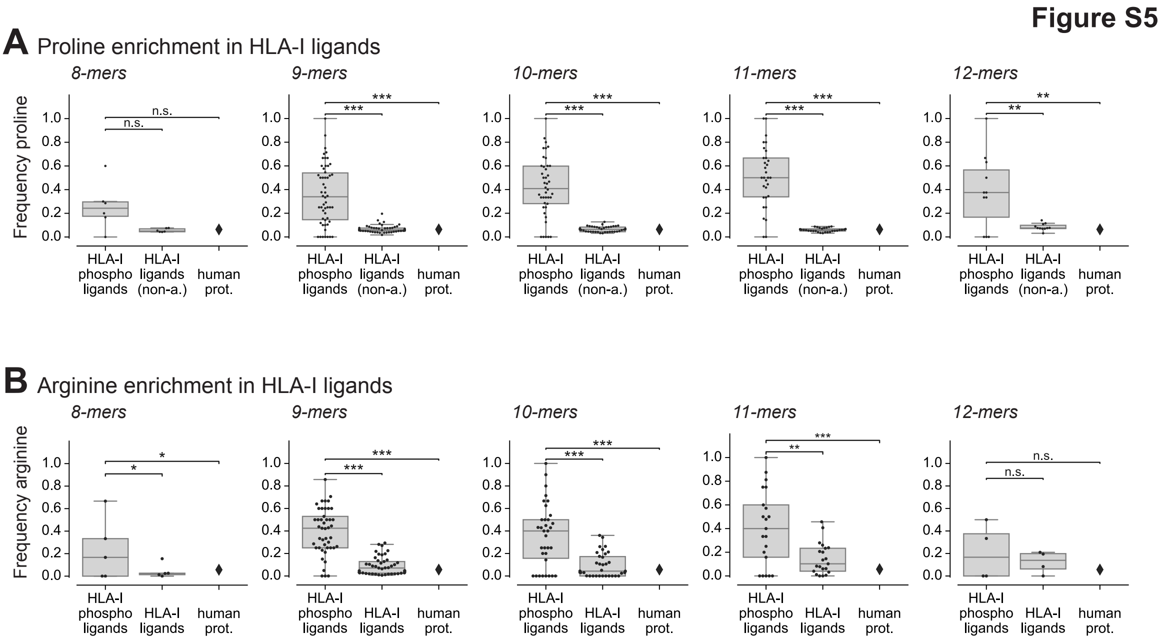


**Figure S5:** Analysis of arginine and proline enrichment for 8- to 12-mers. **(A)** Frequency of proline next to phosphorylated serine in phosphorylated HLA-I peptides (1), proline frequency at non-anchor positions in unmodified HLA-I ligands (2), and proline frequency in the human proteome (3). **(B)** Frequency of arginine at P1 in phosphorylated HLA-I ligands (1), at P1 in unmodified HLA-I ligands (2) and overall arginine frequency in the human proteome (3) for different lengths. (*: $p \le 0.05$; **: $p \le 0.01$; ***: $p \le 0.001$)

Figure S6

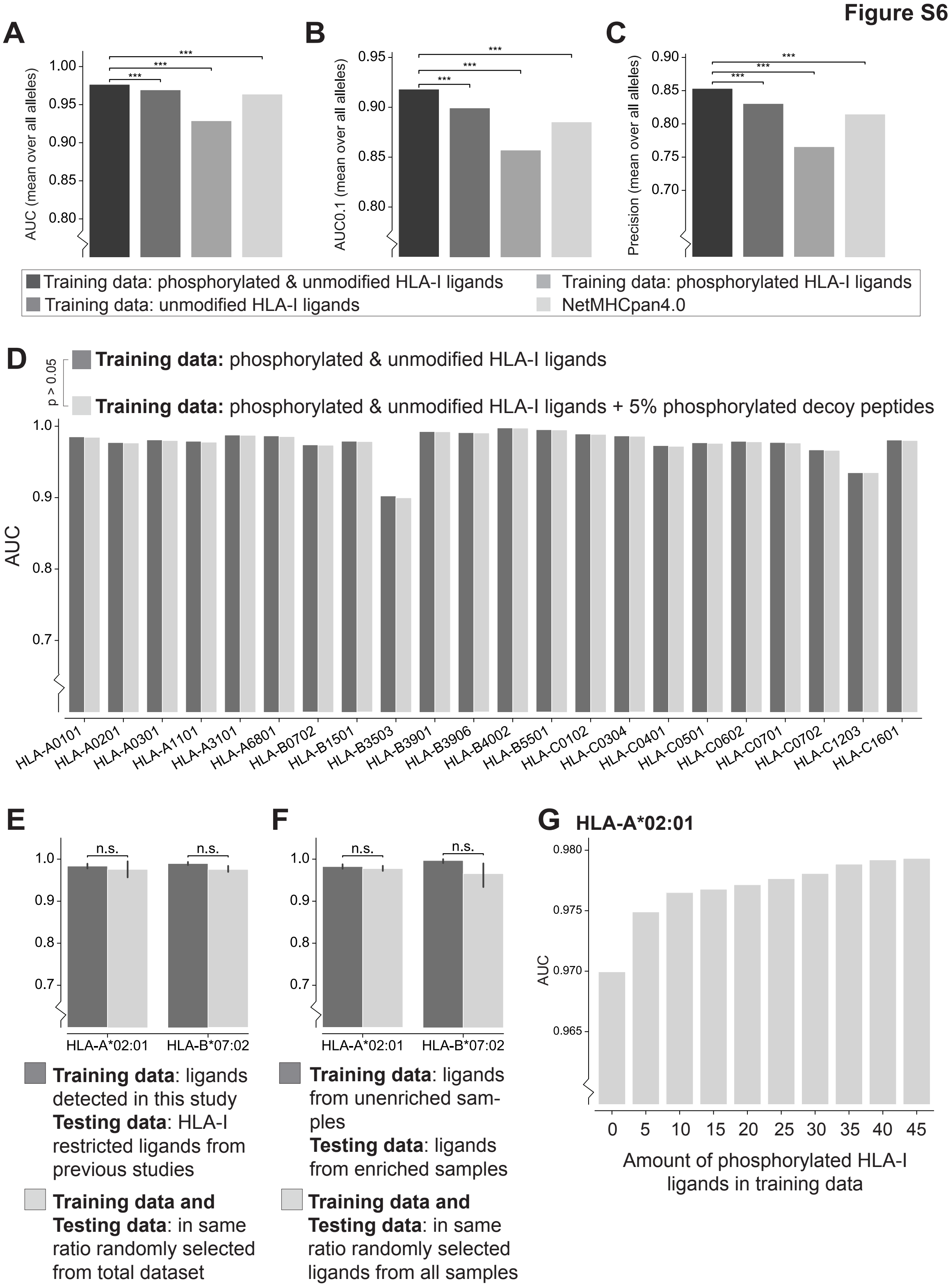


**Figure S6**: 5-fold cross validation of the predictor trained with different training datasets as mean over all alleles shown as **(A)** AUC values, **(B)** AUC0.1, **(C)** Precision of the top 20% of predicted peptides. Training of the predictor was performed with both, phosphorylated and unmodified ligands (1$^{st}$ bar), with only unmodified ligands (2$^{nd}$ bar), and with phosphorylated HLA-I ligands (3$^{rd}$ bar). 4$^{th}$ bar shows AUC values using NetMHCpan4.0 (***: p <= 0.001). **(D)** The predictor was trained on phosphorylated and unmodified HLA-I ligands (left bars); in a second run the predictor was trained on phosphorylated and unmodified HLA-I ligands with additional 5% phosphorylated decoy peptides (right bars). Results are shown as AUC values. **(E)** Predicting previously published phosphorylated HLA-I ligands for HLA-A*02:01 and HLA-B*07:02. The predictor is trained on phosphorylated HLA-I ligands detect by MS in this study. Testing data are HLA-I restricted ligands from previous studies (left bars). As comparison, predictions with random division of training and testing dataset from the combined set of peptides are performed (right bars). **(F)** Prediction of HLA-A*02:01 and HLA-B*07:02. The predictor is trained on phosphorylated HLA-I ligands only found in non-enriched samples and tested on ligands found in enriched samples (left bars). As comparison, predictions with random division of training and testing dataset from the combined set of peptides are performed (right bars). **(G)** Prediction of 10 randomly selected HLA-A*02:01 ligands performed with training data of increasing size.

### Supplementary Tables

**Table S1:** All phosphorylated HLA-I ligands from this study that are known to be phosphorylated by CDK1 or PKA/B from the phosphosite database phosphoELM [3].

**Table S1**

| Kinase | Phosphorylated Peptide | Allele |
|---|---|---|
| CDK1 | VLL[**pS**]PVPEL | HLA-A*02:01 |
| CDK1 | LQL[**pS**]PLKGLSL | HLA-A*02:06, HLA-B*55:01 |
| CDK1 | ITT[**pS**]PITVRK | HLA-A*11:01 |
| CDK1 | EVP[**pT**]PKRPR | HLA-A*68:01 |
| CDK1 | YAS[**pS**]PGGVYATR | HLA-A*68:01 |
| CDK1 | RPI[**pT**]PPRNSA | HLA-B*07:02, HLA-B*55:01 |
| CDK1 | SPK[**pS**]PTAAL | HLA-B*07:02, HLA-B*55:01, HLA-C*03:32 |
| CDK1 | SPRTPV[**pS**]PVKF | HLA-B*07:02 |
| CDK1 | SPR[**pT**]PVSPVKF | HLA-B*07:02 |
| PKA | EPKRR[**pS**]ARL | HLA-B*07:02 |
| PKA | RPRSL[**pS**]SPTV | HLA-B*07:02 |
| PKA | RPRSL[**pS**]SPTVTL | HLA-B*07:02 |
| PKA | RRK[**pS**]HEAEV | HLA-C*06:02 |
| PKB | RAH[**pS**]SPASL | HLA-B*07:02, HLA-B*35:03, HLA-C*01:02, HLA-C*03:03, HLA-C*03:04, HLA-C*03:32, HLA-C*12:03 |
| PKB | RHK[**pS**]DSISL | HLA-B*39:01 |
